# Passive Dynamics Constrain Human Balancing in a Physical Linear-Rail Cart–Pendulum Task

**DOI:** 10.1101/2025.09.27.678964

**Authors:** Laura Alvarez Hidalgo, Ian S. Howard

**Affiliations:** School of Engineering, Computing and Mathematics, University of Plymouth, Plymouth, United Kingdom

**Keywords:** manual balancing, inverted pendulum, motor learning, passive dynamics, sensorimotor control, underactuated systems

## Abstract

Human motor control is often studied using virtual tasks that simplify object mechanics. We examined the stabilization of a physical underactuated cart–pendulum and its relation to passive dynamics. Twelve adults balanced three pendulum configurations (short, 0.31 m; medium, 0.64 m; long, 1.03 m). Free-decay measurements showed progressively lower oscillation frequencies from short to long, while passive-fall durations to ± 30° were longer for the medium and long pendulums than for the short one. Participants trained with the medium pendulum for 30 trials and completed 20 test trials per configuration. Balance duration increased from the first four to the final four training trials, whereas no detectable start-to-end change occurred within any test block. Late-block balance duration was shorter for the short pendulum than for the medium and long pendulums, which did not detectably differ. Estimated peak pendulum angular speed was highest for the short configuration and lowest for the long, whereas peak cart speed did not mirror balance duration. The short configuration, which had the fastest passive dynamics, was balanced for less time than the longer configurations. These results provide a quantitative benchmark for autonomous controllers using a comparable physical linear-rail cart– pendulum under matched mechanical and task constraints.

## Introduction

Humans continually generate and adjust movements to meet changing environmental and task demands. This flexibility depends on both online feedback control and motor learning. Motor learning is often described as involving updates to internal representations of the body and environment, driven in part by sensory prediction errors [1,2]. Such representations may support predictive control, helping the nervous system compensate for sensory noise, feedback delays, and external perturbations.

Much of the experimental understanding of sensorimotor adaptation derives from tightly controlled laboratory tasks using systematic perturbations, such as visuomotor rotations of visual feedback or velocity-dependent curl force fields applied to the hand [3–7]. While these paradigms provide strong experimental control and allow particular learning mechanisms to be isolated [8–10], they typically manipulate selected components of the sensorimotor mapping while holding other properties of the task and apparatus constant. They therefore do not fully capture the interacting effects of mass distribution, inertia, friction, contact forces, and intrinsic instability that arise during the controlling of underactuated physical objects. Important questions consequently remain about how humans learn to stabilize physical objects with different passive dynamics.

Balancing tasks offer an experimentally tractable and ecologically relevant window onto such control problems. They require the timely integration of sensory feedback with anticipatory control to stabilize an intrinsically unstable system. Inverted-pendulum tasks can capture core challenges of physical control, including sensitivity to small disturbances and the destabilizing effects of sensorimotor delays. In a cart–pendulum system, control is also underactuated because the pendulum is not actuated directly but can be influenced only through movement of its base [11,12]. Related challenges arise in postural balance [13,14]; and in the manual balancing of unstable objects [15–17]. Stick-balancing and cart–pendulum paradigms therefore provide useful physical models for investigating how sensorimotor delays and passive mechanical dynamics constrain the time available for corrective action [12,17].

Even studies of unstable motor control have often used robotic interfaces that impose selected dynamics under tightly controlled conditions [18,19]. Simulated inverted-pendulum tasks implemented through robotic manipulanda have enabled controlled manipulation of pendulum dynamics and of the visual, haptic, and temporal feedback available to participants [20–22]. Fully virtual stick-balancing tasks have similarly been used to isolate the effects of feedback delay and the governing dynamical law [23]. By contrast, the present study used a physical cart–pendulum apparatus that provided direct visual observation of the pendulum and direct mechanical interaction with the cart via the handle. Participants stabilized the pendulum by manually moving the cart along a linear rail, thereby controlling the pendulum indirectly through motion of its base.

Linear cart–pendulum systems are canonical benchmark plants in control engineering and have been used to evaluate feedback, optimal, predictive, and learning-based controllers [24–26]. However, studies reporting quantitative human performance together with empirical characterization of the passive dynamics of the same physical pendulum assemblies remain uncommon. Characterizing these passive dynamics alongside training-related changes and balancing performance provides a common reference against which biological and artificial control strategies can subsequently be compared under matched mechanical and task constraints. For otherwise comparable inverted pendulums, increasing effective pendulum length slows the passive dynamics and generally increases the time available for corrective action, making longer pendulums easier to balance than shorter ones.

We tested three pendulum configurations that differed principally in rod length, with additional differences in rod material, rod mass, and assembly-hardware mass. The apparatus was adapted from a cart–pendulum rig developed to investigate engineering approaches to balance control [27–29]. We characterized the passive rotational dynamics of each complete pendulum assembly empirically. The present study had two aims: first, to characterize the passive rotational dynamics of the three configurations using free-decay and passive-fall measurements; and second, to quantify training-related changes and human balancing performance with the same physical apparatus. We examined whether balancing performance differed across configurations in a manner consistent with their measured passive dynamics and predicted that the short configuration would exhibit faster passive dynamics and would be more difficult to balance than the two longer configurations. The paired mechanical and human measurements provide a quantitative reference for subsequent comparisons of biological and autonomous control under comparable task constraints.

## Methods

### AI use statement

The authors used ChatGPT solely as a general research aid for language editing, checking the consistency of manuscript text with author-generated statistical outputs, and identifying potentially relevant literature. ChatGPT was not used to generate data or conduct statistical analyses. All scientific content, analyses, interpretations, and references were independently selected, performed, verified, and approved by the authors.

### Participants

Twelve right-handed adults (8 female, 4 male; mean ± SD age = 24.7 ± 4.0 years) participated in the study. The target sample size of 12 was based on pragmatic recruitment constraints. Participants were randomly assigned to one of two groups (Group A or Group B; *n* = 6 per group) to counterbalance the order of the short- and long-pendulum test conditions. No participants were excluded, and no human-balancing trials were removed from the analysis.

All participants were naïve to the specific experimental hypotheses of the study and provided written informed consent prior to participation. Handedness was confirmed using the Edinburgh Handedness Inventory [30], which verified that all participants were right-handed. The study received ethical approval from the Faculty Research Ethics and Integrity Committee at the University of Plymouth and was conducted in accordance with institutional guidelines.

### Apparatus

The experimental task was implemented using a custom-built inverted-pendulum apparatus, similar to that described in prior work [28,29], mounted on a standard-height desk (Fig. 1). The system consisted of a rigid horizontal frame supporting a rolling cart that moved along the x-axis. A detachable vertical rod was attached to the cart via a pivot joint, forming an underactuated inverted pendulum whose angle could be controlled only indirectly through movement of the cart.

**Figure 1.**
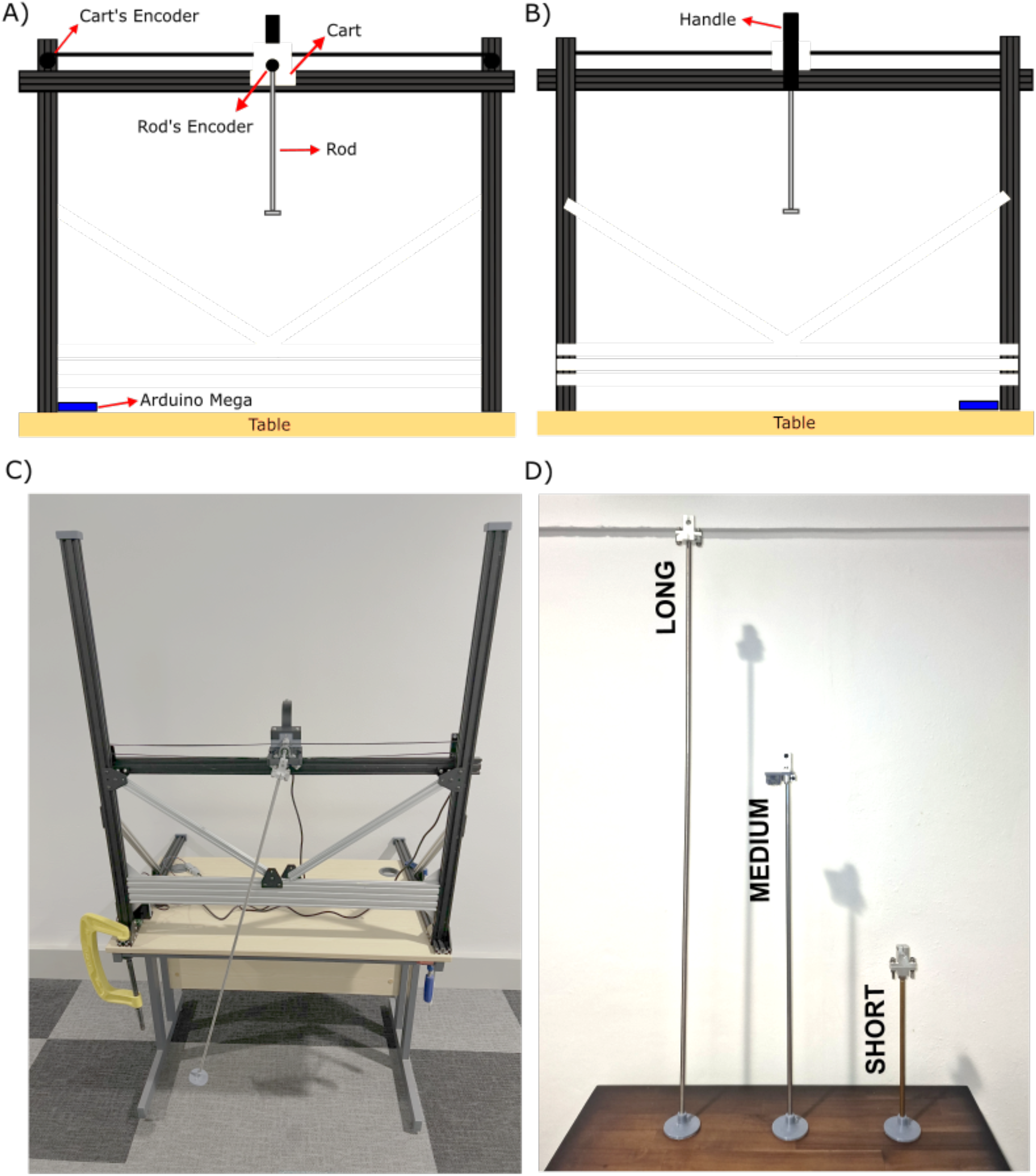
Schematic and photographic views of the inverted pendulum apparatus. **A)** Front view of the setup, showing the rolling cart and vertical rod assembly. **B)** Rear view, highlighting the handle used for manual control. **C)** Photograph of the complete rig mounted on a standard-height desk. **D)** The three interchangeable pendulum rods (short, medium, and long).

Each rotary encoder provided 2400 counts per revolution after quadrature decoding, corresponding to an encoder-shaft angular resolution of 0.15° per count. One encoder was mounted at one end of the rail and measured horizontal cart displacement through a GT2 timing belt attached to the cart and a 36-tooth GT2 pulley. This transmission produced 72 mm of belt travel per pulley revolution, corresponding to a linear cart-position resolution of 0.030 mm per encoder count. The second encoder was located at the pendulum–cart pivot and measured pendulum angle.

Three interchangeable 8-mm-diameter rods were used to vary the passive dynamics of the pendulum. The short assembly used a 0.305 m brass rod, whereas the medium and long assemblies used 0.635 m and 1.030 m stainless-steel rods, respectively. The rods shared a similar mounting interface to maintain consistent mechanical coupling with the cart.

Each complete moving pendulum assembly comprised the rod, an aluminum pivot-mount attachment, an end cap, and associated fasteners. The complete assemblies were weighed in their assembled configuration, and the component masses reported in Table 1 were determined separately. Because the complete assemblies contained non-rod components with a nonuniform spatial distribution, they were not treated as uniform rods. Instead, the effective inertia-normalized dynamical coefficients of each complete pendulum assembly were estimated directly from the measured free-decay trajectories.

**Table 1.** Measured physical properties of the three pendulum assemblies. Rod length and bare rod mass were measured for each interchangeable pendulum. Total pendulum assembly mass comprised the bare rod, pivot-mount attachment, end cap, and associated fasteners. The moving cart mass and usable cart travel were common to all configurations.

| Property | Short pendulum | Medium pendulum | Long pendulum |
| --- | --- | --- | --- |
| Rod material | Brass | Stainless steel | Stainless steel |
| Rod length, $L$ (m) | 0.305 | 0.635 | 1.030 |
| Bare rod mass, $m_r$ (kg) | 0.117 | 0.236 | 0.397 |
| Attachment mass, $m_a$ (kg) | 0.053 | 0.065 | 0.055 |
| Endcap mass, $m_c$ (kg) | 0.021 | | |
| Total pendulum assembly mass, $m_{tot}$ (kg) | 0.191 | 0.322 | 0.473 |
| Cart mass, excluding the pendulum assembly (kg) | 0.767 |  |  |
| Usable cart travel (m) | 0.785 |  |  |

Rod length was the principal design difference, but material, rod mass, and combined assembly-hardware mass also differed across assemblies. The conditions are therefore referred to as short, medium, and long pendulum assemblies rather than as a pure manipulation of length.

Participants controlled the cart through a rear-mounted handle. The entire rig was securely clamped to a standard-height desk to maintain mechanical stability. An Arduino Mega 2560 microcontroller recorded pendulum angle and cart position at an effective sampling frequency of approximately 6 Hz, with each sample stored together with its timestamp.

### Experimental Setup

Two complementary studies used the same inverted-pendulum apparatus (Fig. 1). Study 1 characterized the passive fixed-pivot rotational behavior of the three pendulum assemblies without human input. Study 2 examined human balancing performance using the same assemblies. A ± 30° threshold was used for the passive-fall and human-balancing trials; the free-decay recordings continued without an angular termination threshold. Each recording or trial began with the cart positioned approximately at the center of the rail. The offset of a 3 s start tone cued the beginning of data acquisition, and a second 3 s tone signaled its end.

### Study 1: Passive Pendulum Characterization

To characterize the passive dynamics of the three pendulum assemblies, measurements were obtained without human control using two protocols: a continuous free-decay recording and repeated passive falls from upright to the ± 30° threshold.

For the passive free-decay recordings, the researcher raised the pendulum from its stable downward position towards the unstable upright position and held it stationary. Once the absolute encoder-reported angular displacement reached at least 180° from its initial position and its online finite-difference estimate of angular speed was no greater than 5° s^−1^, a 3 s, 440 Hz start tone was presented. The researcher continued to hold the pendulum upright during the tone. At tone offset, the acquisition program stored the current pendulum and cart encoder counts as zero references and began data acquisition.

Tone offset simultaneously cued the researcher to release the pendulum without further contact. The resulting free motion was recorded continuously for approximately 100–300 s, depending on pendulum length, until the oscillations had substantially decayed. A second 3 s, 440 Hz tone signaled the end of data acquisition. One free-decay recording was obtained for each pendulum assembly. For model fitting, the recorded angle was subsequently re-referenced to the empirically estimated stable downward equilibrium using the median-based procedure described below, and only the retained oscillatory portion of the trajectory was analyzed.

The passive-fall trials used the same positioning, cueing, zero-referencing, and release procedure as the free-decay recordings. Each trial ended when the absolute angular displacement first reached 30°, or after 10 s if the threshold had not been reached. Passive-fall duration was defined as the elapsed time from the software-defined start of data acquisition at tone offset to the first threshold crossing, capped at 10 s.

Twenty passive-fall trials were planned for each pendulum assembly. All 20 trials were available for the short and long assemblies. Owing to a recording error, only 19 valid trials were available for the medium assembly. Because the releases were performed manually, some variability in the physical release angle and angular velocity was unavoidable.

### Study 2: Human Study of Balance

Before the experiment, participants were instructed to control the cart by moving its handle horizontally while maintaining their gaze near the upper end of the pendulum. They were instructed to keep the cart stationary during the start tone, begin controlling the cart when the tone ended, and stop moving when the end tone was presented.

During each trial, the participant held the cart handle while the researcher slowly raised the pendulum from its downward resting position to the unstable upright position. The acquisition program monitored pendulum angle and an online finite-difference estimate of angular velocity.

Once the absolute encoder-reported angular displacement reached at least 180° from its initial position and its estimated angular speed was no greater than 5° s^−1^, a 3 s, 440-Hz start tone was presented. The researcher continued to hold the pendulum upright, and the participant kept the cart stationary, throughout the tone.

At tone offset, the acquisition program stored the current pendulum and cart encoder counts as zero references and started data acquisition. Tone offset cued the researcher to release the pendulum and the participant to begin controlling the cart. Thus, trial time *t* = 0 corresponded to the software-defined start of data acquisition, and *ϕ* = 0° represented the upright pendulum position immediately before release.

The trial ended when the absolute pendulum angle first reached 30°, or after 10 s if the participant maintained the pendulum within the permitted angular range. Balance duration was the elapsed time from the start of data acquisition to the first threshold crossing, capped at 10 s.

A second 3 s, 440 Hz tone was presented immediately after data acquisition ended, signaling trial completion to both the participant and the researcher.

Participants received no augmented performance feedback. They could observe the pendulum and experience the duration and outcome of each trial but were not shown numerical performance measures. No additional scheduled rest breaks were provided beyond the brief intertrial intervals, although participants rested while the pendulum assemblies were changed.

Each participant completed one training block followed by three testing blocks (Fig. 2). During the training block, participants always used the medium pendulum assembly and completed 30 trials. They then completed a 20-trial test block with the medium assembly, followed by 20-trial test blocks with the short and long assemblies in counterbalanced order. Participants therefore completed one of two testing orders: medium–short–long or medium–long–short.

**Figure 2.**
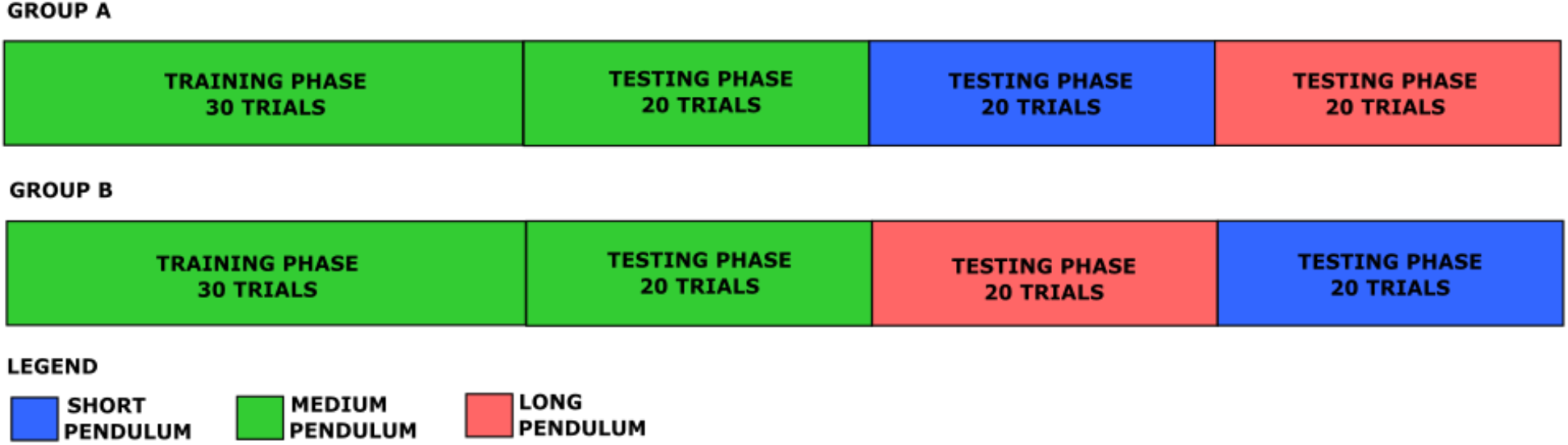
Experimental design and trial structure across conditions. Each row shows the sequence of phases for one group. All participants trained with the medium assembly for 30 trials, then completed 20-trial test blocks with the medium assembly first, followed by the short and long assemblies in counterbalanced order. Thus, one group completed the order medium–short–long, whereas the other completed medium–long–short. Green, blue, and red denote the medium, short, and long assemblies, respectively.

Because the medium assembly served as both the training condition and the first test condition, comparisons involving it were not fully counterbalanced.

Each testing block comprised 20 consecutive trials using one pendulum assembly. In total, each participant completed 90 trials: 30 during training and 60 during testing. The experiment lasted approximately 60 min per participant.

Participants performed the task while standing and controlled the cart using their right hand. Contact with a rail limit did not itself terminate a trial.

### Pendulum Measurements

The rotary encoders provided time-indexed measurements of horizontal cart position and pendulum angle. Cart position, *x*(*t*), was calculated from the cart-encoder count by converting pulley rotation into linear displacement through the GT2 timing-belt transmission.

During the human-balancing and passive-fall trials, pendulum angle, *ϕ*(*t*), was expressed relative to the unstable upright equilibrium, such that

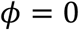

corresponded to the vertically upright pendulum. The angular termination threshold was reached when

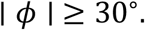

For the free-decay analysis, the recorded pendulum angle was re-referenced to the stable downward equilibrium. This angle is denoted by *θ*(*t*), with

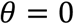

corresponding to the downward resting position.

### Passive rotational characterization of the pendulum assemblies

The single free-decay recording obtained for each assembly was used for passive rotational characterization, with the original measurement timestamps retained for model fitting.

For each assembly, the maximum absolute cart displacement during the retained fitting interval was one encoder count, equivalent to 0.03 mm, and was therefore considered negligible. The rod-and-mount assembly was therefore modeled as a compound physical pendulum rotating about a fixed pivot. For model fitting and numerical integration, pendulum angle and angular velocity were expressed in radians and radians per second, respectively. For rotating components *i*, with mass *m*_*i*_, center-of-mass distance *r*_*i*_ from the pivot, and center-of-mass moment of inertia *I*_*i*,COM_ the effective rotational inertia was

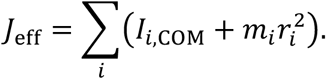

The passive equation of motion was

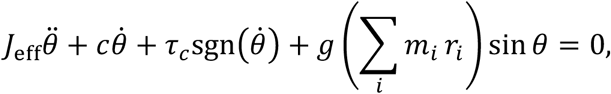

where *c* is the rotational viscous-damping coefficient, *τ_c_* is the magnitude of the Coulomb-friction torque, and *g* = 9.81 m s^−2^. Dividing by *J*_*eff*_ and replacing sgn 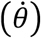 with the smooth approximation tanh 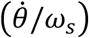 for numerical fitting gave the inertia-normalized model

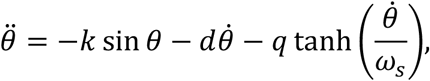

with

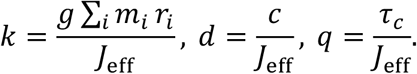

The hyperbolic-tangent term provided a smooth approximation to Coulomb friction near angular-velocity reversals. The transition velocity was fixed at

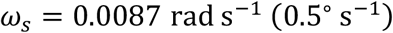

for all three pendulums.

The terminal settled level was estimated as the median of the final 20 angular samples. The final sample whose absolute deviation from this level exceeded 3 × 10^−4^ rad was defined as the last moving sample, and all subsequent samples were excluded because they corresponded to the terminal settled or sticking region, and static friction was not explicitly represented by the model. The remaining angular record was provisionally expressed relative to the nominal stable downward position. Positive angular peaks were identified using MATLAB’s findpeaks function, and the first peak with an angular displacement no greater than 30° was selected as the beginning of the fitting interval. The retained fitting interval extended from this peak to the final retained moving sample.

This task-defined cutoff used the same angular magnitude as the ± 30° termination threshold in the passive-fall and human-balancing trials, although the angles were referenced to different equilibria. It restricted parameter estimation to the moderate- and low-amplitude regime most relevant to the experimental task and reduced the influence of unmodeled high-speed effects during the initial swings.

Oscillation frequency was estimated from the first 11 positive angular peaks within the retained fitting interval whose amplitudes were less than or equal to 30°. The oscillation period was calculated as the mean of the 10 intervals between successive peaks, and oscillation frequency was calculated as the reciprocal of this mean period.

A constant equilibrium offset was then calculated separately for each assembly as the median of all raw angular samples within the retained fitting interval. Denoting the retained interval by *J*, the offset was

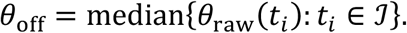

The offset-corrected measured angle was calculated as

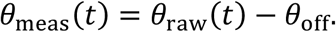

The resulting angular record was therefore centered on the empirically estimated stable downward equilibrium, and no angular offset parameter was included in the model optimization.

The selected turning point was defined as *t* = 0, and the model was initialized using

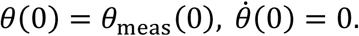

Beginning at a turning point reduced sensitivity to uncertainty in the manually generated release angle and velocity.

Initial conditions and parameter bounds for *k, d*, and *q* are shown in Supplementary Table S1. The optimization vector was

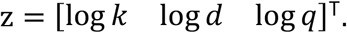

The three positive dynamical coefficients were represented in logarithmic coordinates to enforce positivity during optimization, improve numerical scaling across coefficients of different magnitudes, and allow multistart perturbations to act proportionally rather than additively. The smoothing velocity, *ω*_*S*_, was fixed rather than estimated.

The parameters *k, d*, and *q* were estimated by bounded nonlinear least squares using MATLAB’s lsqnonlin function. At each objective-function evaluation, the model was integrated using ode45 and evaluated at the original measurement times. Up to five fitting attempts were performed. The first used the condition-specific initial estimate, and subsequent attempts used random local perturbations of *k, d*, and *q* in logarithmic parameter space, with a standard deviation of 0.7. Perturbed starting points were restricted to the specified parameter bounds. The multistart sequence was terminated when a completed fit achieved an angular RMSE of 0.5° or less; otherwise, all five starts were attempted. Among the successfully completed attempts, the solution with the smallest residual sum of squares was retained, irrespective of the optimization exit flag. Numerical optimization settings are reported in Supplementary Table S2.

Fit quality was calculated directly from the difference between the fitted model trajectory and the offset-corrected measured trajectory. Angular root-mean-square error was calculated as

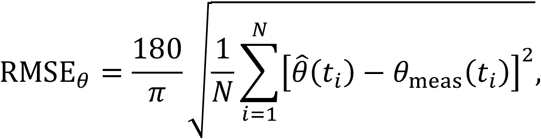

expressed in degrees. The fitted and measured trajectories were also inspected to verify agreement in the oscillation phase and the amplitude-decay envelope. The estimated coefficients were interpreted as effective descriptive parameters of the complete rod-and-mount assemblies.

For an independent physical consistency check, a nominal gravitational coefficient was calculated for a uniform bare rod pivoted at one end:

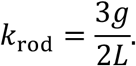

These nominal values were calculated from measured rod length and were not imposed during model fitting.

### Angle-based kinematics

To characterize system kinematics, pendulum angle and angular velocity, together with cart position and linear velocity, were calculated. For the descriptive kinematic analyses, pendulum angle was expressed in degrees, pendulum angular velocity in degrees per second, cart position in meters, and cart velocity in meters per second.

During acquisition, an online finite-difference estimate of pendulum angular velocity was calculated in MATLAB from successive encoder readings and used to determine whether the pendulum was sufficiently stationary before each recording or trial. For offline analysis, pendulum encoder counts were converted to degrees and corrected for angular wraparound, while cart encoder counts were converted to meters using the GT2 timing-belt transmission.

The recorded cart-position and pendulum-angle signals were interpolated onto a regular 100-Hz time vector using the MATLAB interp1 function with the spline option. Approximate cart velocity and pendulum angular velocity were then calculated using the MATLAB gradient function with a sampling interval of 0.01 s. The interpolated angular trajectory was also used to estimate the time at which the pendulum crossed the ± 30° failure threshold. Because the original position signals were acquired at an effective sampling frequency of approximately 6 Hz, the derived speed measures were treated as approximate descriptive estimates rather than high-precision kinematic measurements.

### Outcome measures

Balance duration was defined as the elapsed time from the start of data acquisition until the absolute pendulum angle relative to upright first reached or exceeded 30°. Trials were capped at 10 s; when no threshold crossing occurred, balance duration was recorded as 10 s.

Estimated peak cart speed was defined as the maximum absolute estimated cart velocity within a trial. Estimated peak pendulum angular speed was defined as the maximum absolute estimated pendulum angular velocity within a trial.

### Data and statistical analysis

Experimental data were processed offline using MATLAB R2026a (The MathWorks, Natick, MA, USA). Statistical analyses were conducted using JASP 0.19.1 (JASP Team, Amsterdam, the Netherlands).

For each experimental block, participant-level balance duration was averaged separately across the first four and final four trials. The four-trial averaging windows were selected to reduce trial-level variability while retaining sensitivity to early- and late-block performance. These values were analyzed using a mixed repeated-measures ANOVA with Block (medium training, medium test, short test, and long test) and Phase (start and end) as within-participant factors and short–long presentation order (medium–short–long or medium–long–short) as a between-participant factor.

Late-block balance duration was compared across the three test conditions using a one-way repeated-measures ANOVA applied to participant-level means from the final four trials of each test block. A separate Condition × Order mixed ANOVA, with Condition as a within-participant factor and presentation order as a between-participant factor, was used to examine whether short–long presentation order influenced the late-block condition comparison.

Participant-level estimated peak cart speed and peak pendulum angular speed were also averaged across the final four trials of each test condition and analyzed separately using mixed repeated-measures ANOVAs, with Condition as a within-participant factor and presentation order as a between-participant factor.

For within-participant effects containing more than two levels, sphericity was assessed using Mauchly’s test. When the assumption of sphericity was violated, Greenhouse– Geisser-adjusted degrees of freedom and p-values were reported. For each ANOVA, follow-up pairwise comparisons were adjusted using Holm’s procedure within that family of comparisons. All tests were two-sided, with the significance level set at *α* = 0.05. Effect sizes were reported as omega squared 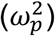.

Passive-fall durations were summarized descriptively using the mean and standard deviation. Because these repeated releases were obtained from a single apparatus and operator, they were not treated as independent population-level replicates, and no inferential statistical tests were applied to the passive-fall observations.

## Results

### Study 1: Passive Pendulum Characterization

#### Free Decay

The retained free-decay trajectories showed progressively lower oscillation frequencies across the short, medium, and long pendulum assemblies. The estimated oscillation frequency was 1.06 Hz for the short assembly, 0.74 Hz for the medium assembly, and 0.58 Hz for the long assembly.

Figure 3 panels A-C show the measured and fitted free-decay trajectories over the retained fitting intervals for the three pendulum assemblies. For visualization in Figure 3, the measured trajectories were interpolated to 100 Hz; model fitting was performed at the original measurement timestamps.

**Figure 3.**
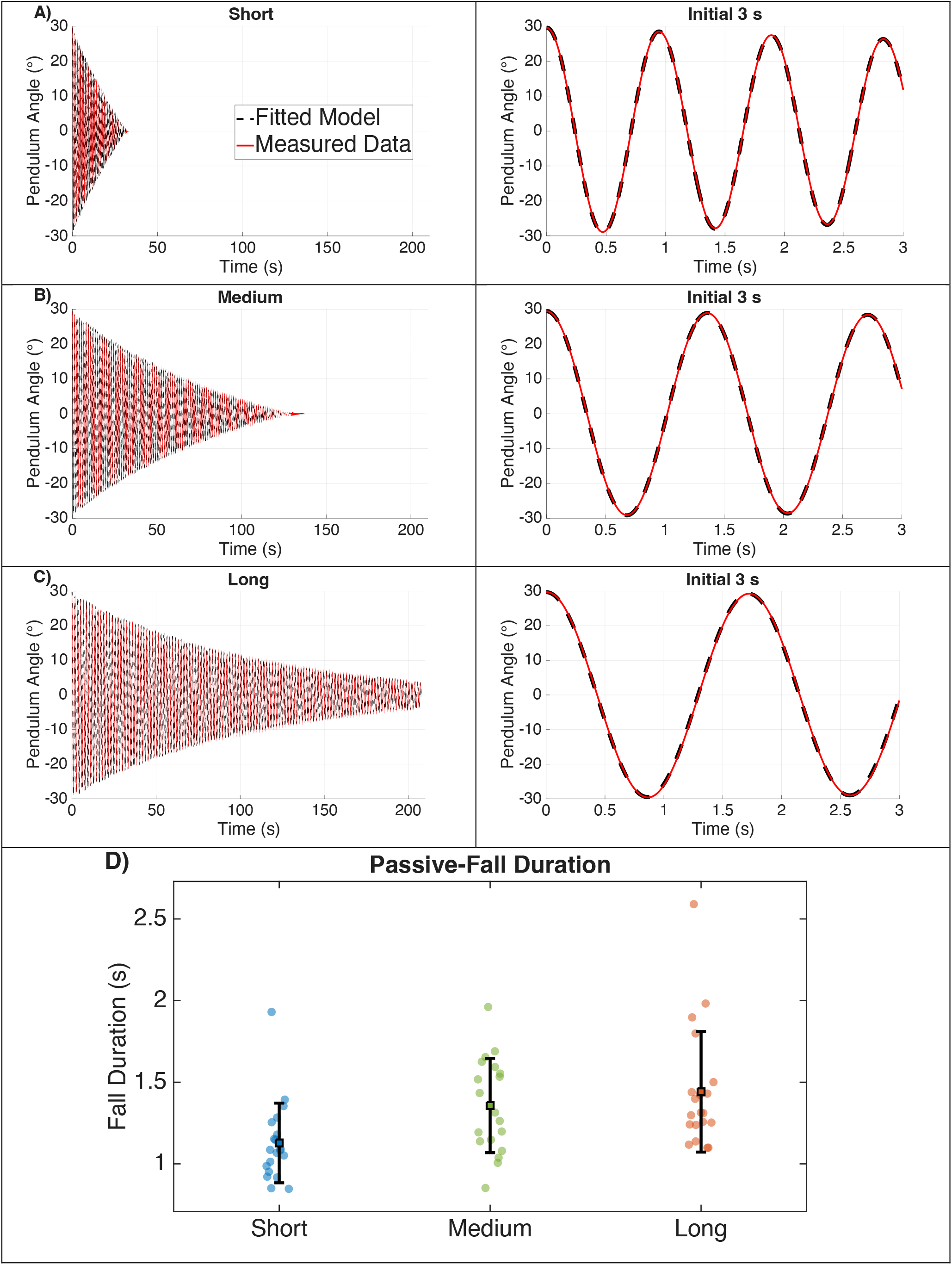
Passive characterization of the three pendulum assemblies. A–C) Measured and fitted free-decay trajectories for the short (A), medium (B), and long (C) pendulum assemblies. Left panels show the complete retained fitting intervals on a common time scale; right panels show the first 3 s on an expanded scale. Measured, offset-corrected pendulum angles are shown as red solid lines and fitted model trajectories as black dashed lines. Time zero denotes the first positive angular peak identified using MATLAB’s findpeaks function for which the displacement from the stable downward equilibrium was no greater than 30°. D) Passive-fall duration to the ± 30° threshold for the short (*n* = 20), medium (*n* = 19), and long (*n* = 20) assemblies. Individual releases are shown with horizontal jitter. Black squares indicate condition means, and error bars denote ± 1 SD.

The fitted models reproduced the oscillation phase and amplitude-decay envelope of the retained free-decay trajectories, with angular RMSEs of 0.293°, 0.365°, and 0.275° for the short, medium, and long assemblies, respectively. Fitted and nominal coefficients are reported in Table 2.

**Table 2.**
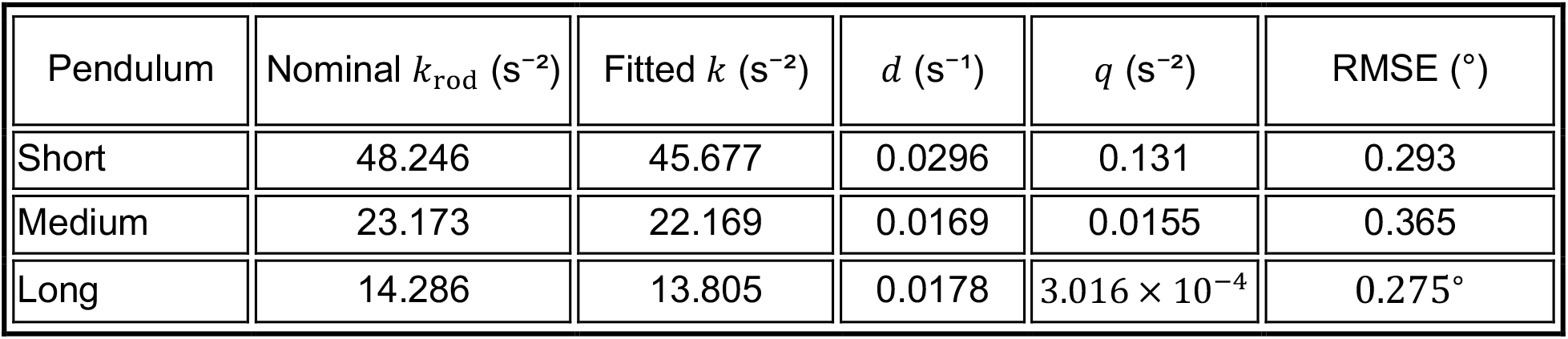
Inertia-normalized coefficients estimated from the free-decay trajectories and comparison with nominal uniform-rod gravitational coefficients. Nominal values were calculated as *k*_*rod*_ = 3*g*/(2*L*), assuming a uniform rod pivoted at one end, and were not imposed during fitting. The smoothing velocity was fixed at *ω*_*s*_ = 0.0087 rad *s*^−1^ for all three pendulums. RMSE denotes angular root-mean-square error over the retained fitting interval.

| Pendulum | Nominal $k_{\text{rod}}$ ($\text{s}^{-2}$) | Fitted $k$ ($\text{s}^{-2}$) | $d$ ($\text{s}^{-1}$) | $q$ ($\text{s}^{-2}$) | RMSE ($^\circ$) |
| --- | --- | --- | --- | --- | --- |
| Short | 48.246 | 45.677 | 0.0296 | 0.131 | 0.293 |
| Medium | 23.173 | 22.169 | 0.0169 | 0.0155 | 0.365 |
| Long | 14.286 | 13.805 | 0.0178 | $3.016 \times 10^{-4}$ | $0.275^\circ$ |

For the long pendulum, the fitted Coulomb-like coefficient was *q* = 3.016 × 10^−4^ s^−2^, substantially smaller than the corresponding coefficients for the short and medium pendulums and toward the lower end of the permitted parameter range. The Coulomb-like contribution was therefore treated as effectively negligible or weakly resolved.

#### Passive falls to ±30°

Passive-fall duration (mean ± SD) was 1.128 ± 0.244 s for the short assembly (*n* = 20), 1.357 ± 0.289 s for the medium assembly (*n* = 19), and 1.441 ± 0.369 s for the long assembly (*n* = 20). One planned medium-assembly observation was unavailable because of a recording error. Within this apparatus and manual-release procedure, the short assembly reached the ± 30° threshold more quickly than the medium and long assemblies. Figure 3D shows the passive fall durations.

### Study 2: Human Study of Balance

#### Balance Duration Across Training and Testing Blocks

As shown in Fig. 4, balance duration increased from the beginning to the end of the initial medium-pendulum training block. The prespecified Block × Phase analysis compared participant-level means from the first four and final four trials of each block.

**Figure 4.**
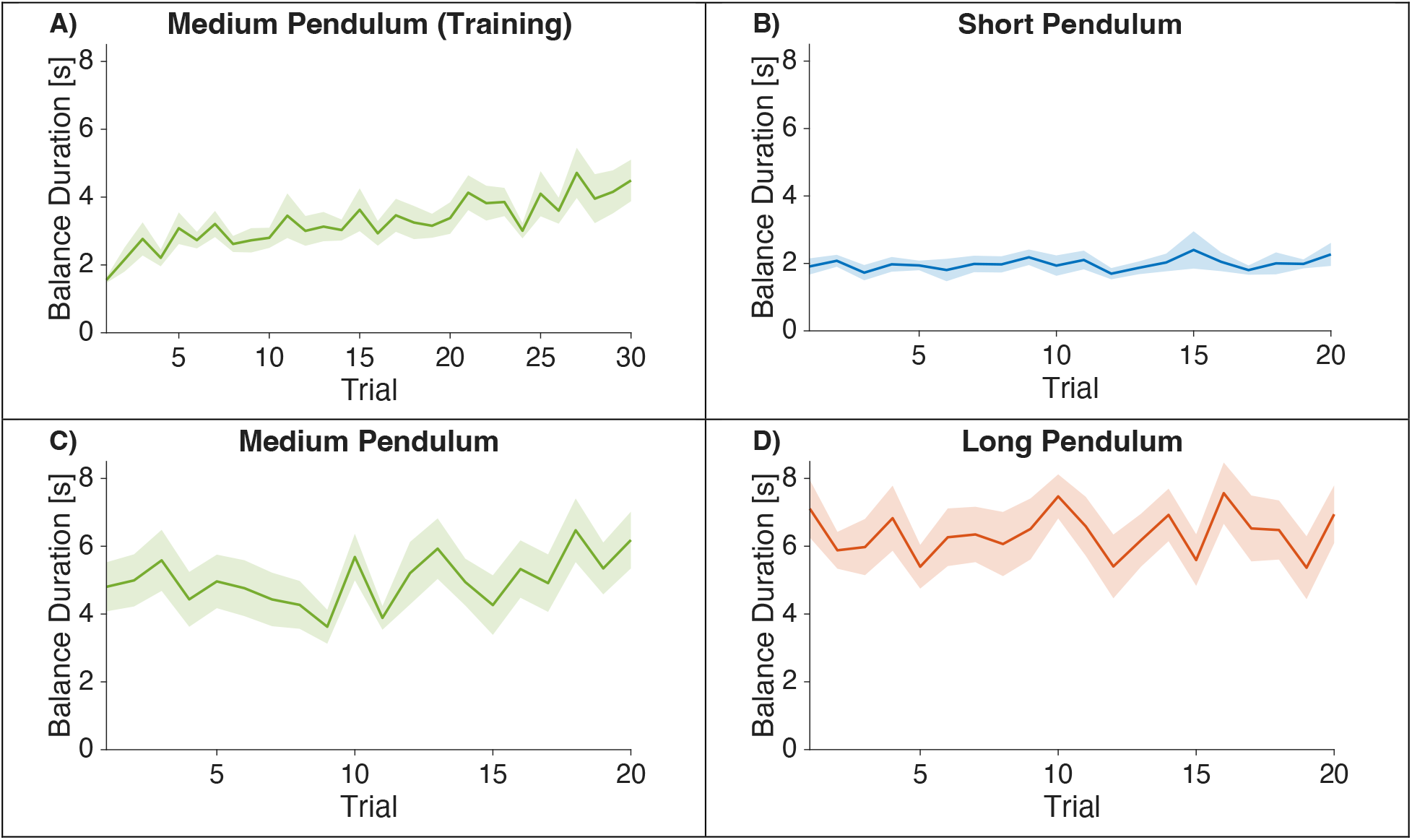
Balance duration across training and testing trials. A) Medium-pendulum training block. B) Short-pendulum test block. C) Medium-pendulum test block. D) Long-pendulum test block. Lines show group means, and shaded bands denote ± 1 SE across participants.

**Figure 5.**
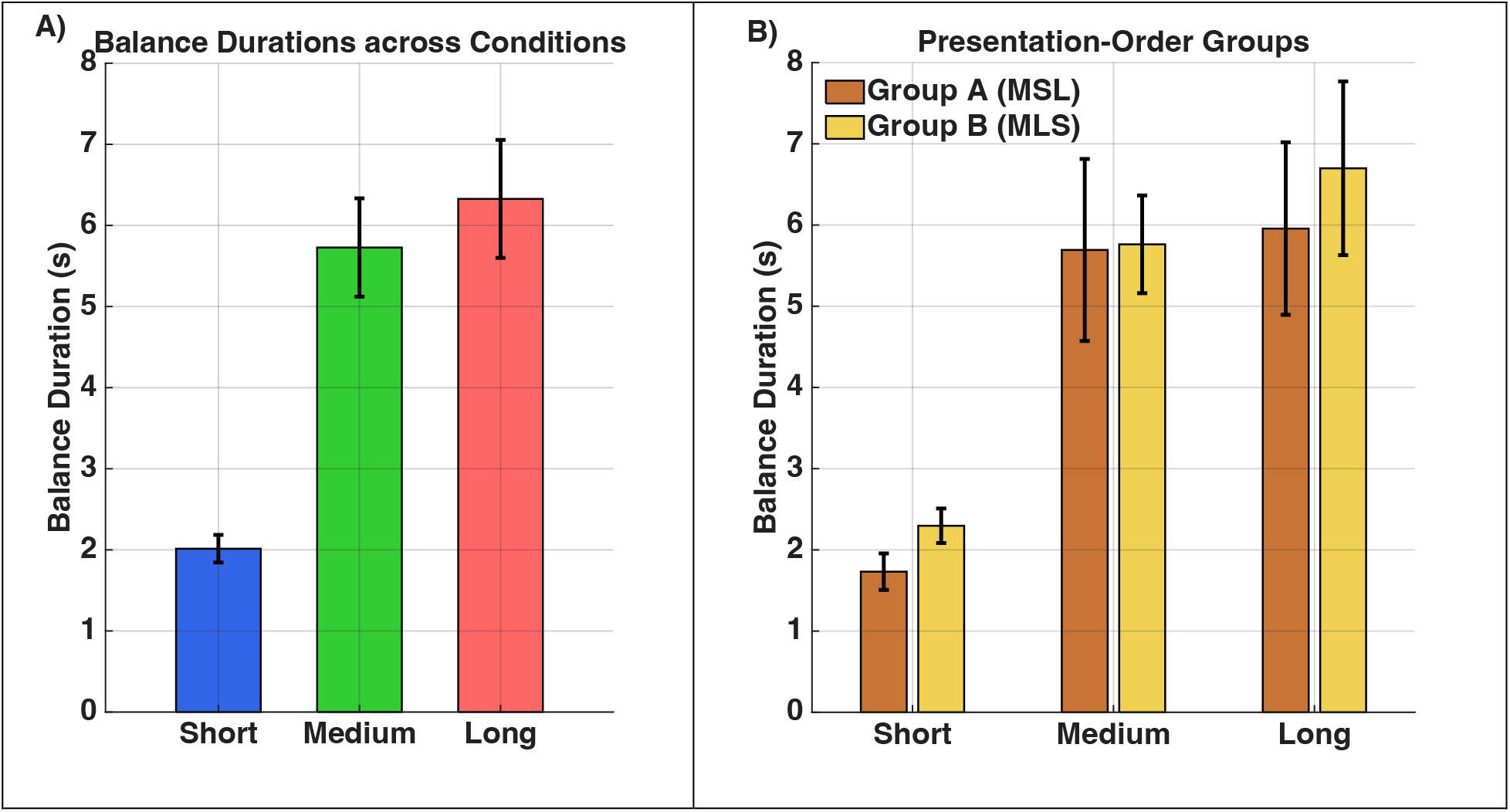
Late-block balance duration averaged across the final four trials. Error bars denote ±1 SE across participants. **A)** Condition means for the short, medium, and long assemblies. B) Condition means shown separately for the medium–short–long and medium–long–short presentation-order groups.

There were significant main effects of Block, *F*(3, 30) = 47.652, *p* < 0.001, *ω*^2^ = 0.551, and Phase, *F*(1, 10) = 15.419, *p* = 0.003, *ω*^2^ = 0.064, together with a significant Block × Phase interaction, *F*(3, 30) = 5.748, *p* = 0.003, *ω*^2^ = 0.073. No statistically detectable effects involving presentation order were observed (all *p* ≥ 0.298). Mauchly’s tests did not indicate violations of sphericity for Block, *W* = 0.293, *p* = 0.059, or the Block × Phase interaction, *W* = 0.437, *p* = 0.207.

Balance duration (mean ± SD across participants) increased from 2.173 ± 0.778 s during the first four medium-training trials to 4.328 ± 1.827 s during the final four trials (*p* = 0.009). No statistically detectable start-to-end changes were observed in the short test block (1.922 ± 0.515 to 2.014 ± 0.590 s, *p* = 1.000), medium test block (4.953 ± 2.256 to 5.728 ± 2.101 s, *p* = 0.462), or long test block (6.447 ± 2.055 to 6.328 ± 2.519 s, *p* = 1.00). Pairwise *p*-values were adjusted using Holm’s procedure.

#### Late-Block Balance Duration Across Pendulum Conditions

The only statistically detectable start-to-end increase occurred during the medium training block. Late-block balance duration was compared using the mean of the final four trials from each testing block. Mean late-block balance duration was 2.014 ± 0.590 s for the short assembly, 5.728 ± 2.101 s for the medium assembly, and 6.328 ± 2.519 s for the long assembly. A one-way repeated-measures ANOVA showed a significant effect of condition, *F*(2, 22) = 33.136, *p* < 0.001, *ω*^2^ = 0.488. Holm-adjusted pairwise comparisons showed that late-block balance duration was shorter for the short assembly than for the medium and long assemblies (both *p* < 0.001); the medium and long assemblies did not differ detectably (*p* = 0.246). Mauchly’s test did not indicate a violation of sphericity, *W* = 0.825, *p* = 0.383.

In a separate mixed ANOVA including short–long presentation order, neither the main effect of Order, *F*(1, 10) = 0.247, *p* = 0.630, *ω*^2^ = 0.000, nor the Condition × Order interaction, *F*(2, 20) = 0.171, *p* = 0.844, *ω*^2^ = 0.000, was statistically detectable. The medium condition was not counterbalanced because it served as the trained reference.

The 10-s task limit was not reached in any of the 48 late-block short-pendulum trials (the final four trials for each of the 12 participants), but was reached in 9 of the 48 medium-pendulum trials (18.8%) and 14 of the 48 long-pendulum trials (29.2%).

#### Estimated Peak Cart and Pendulum Speeds

To describe movement kinematics across pendulum conditions, estimated peak cart speed and pendulum angular speed were compared over the final four trials. Because these estimates were derived from low-frequency position records, they were treated as approximate kinematic estimates rather than precise kinematic measurements.

As shown in Fig. 6C, estimated peak cart speed over the final four trials differed across pendulum conditions. Mean estimated peak cart speed was 0.911 ± 0.173 m s^−1^ in the short condition, 1.062 ± 0.330 m s^−1^ in the medium condition, and 0.810 ± 0.224 m s^−1^ in the long condition. Mean estimated peak cart speed was therefore numerically highest in the medium condition and lowest in the long condition. Mauchly’s test indicated a violation of sphericity, *W* = 0.492, *x*^2^(2) = 6.384, *p* = 0.041; therefore, Greenhouse–Geisser-corrected degrees of freedom were reported, with *ε*_GG_ = 0.663. The mixed repeated-measures ANOVA showed a significant main effect of pendulum condition on estimated peak cart speed, *F*(1.326, 13.262) = 9.301, *p* = 0.006, *ω*^2^ = 0.156. There was no significant main effect of presentation order, *F*(1, 10) = 2.883, *p* = 0.120, *ω*^2^ = 0.079, and no significant Condition × Order interaction, *F*(1.326, 13.262) = 0.310, *p* = 0.650, *ω*^2^ < 0.001. Holm-adjusted pairwise comparisons, averaged across presentation orders, showed that estimated peak cart speed was significantly higher in the medium than in the long condition (*p* = 0.005) and significantly higher in the short than in the long condition (*p* = 0.039); the short and medium conditions did not differ significantly (*p* = 0.071).

**Figure 6.**
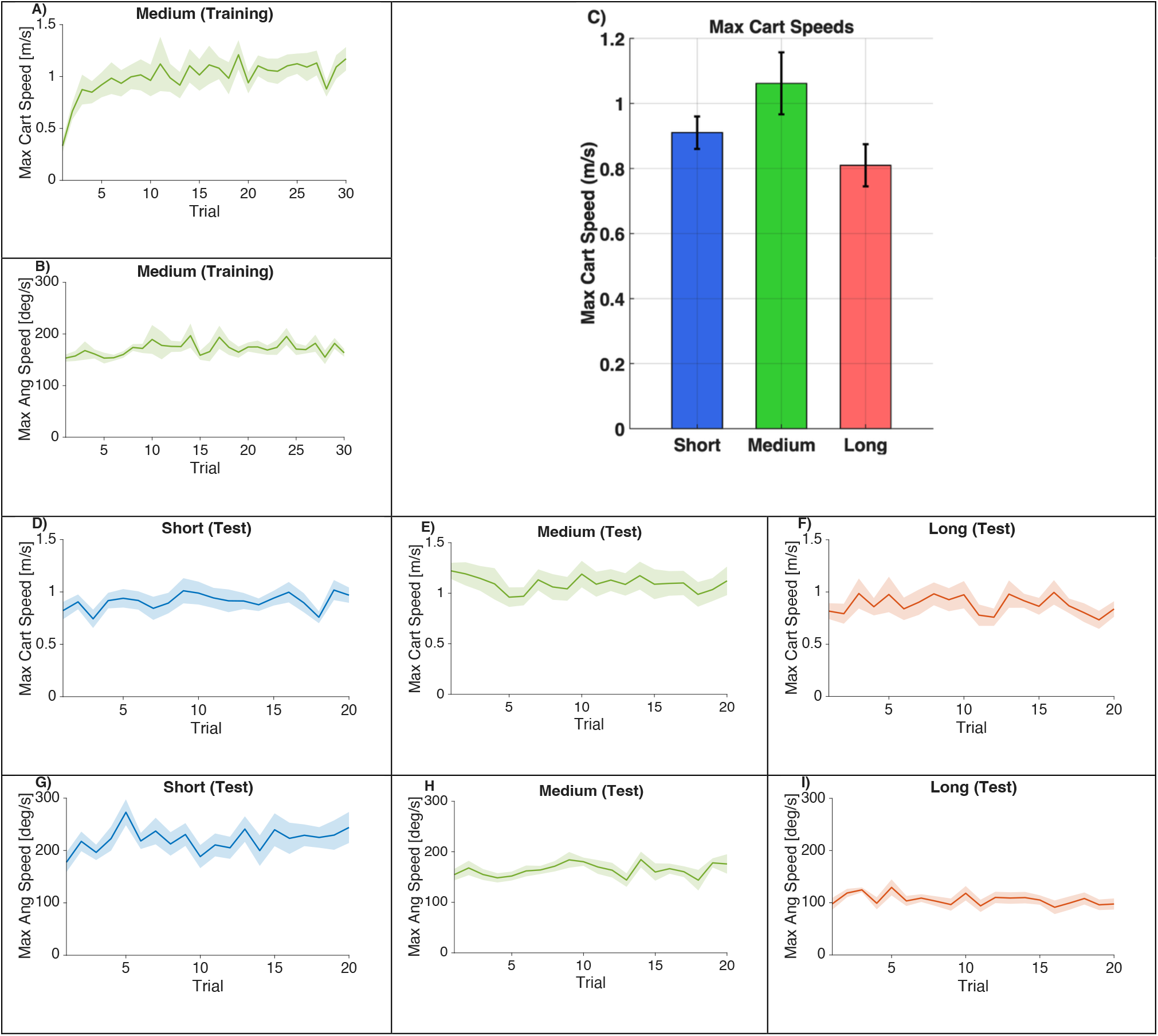
Estimated peak speeds across pendulum conditions and experimental phases. A–B) Per-trial estimated peak cart speed (A) and peak pendulum angular speed (B) across the 30 medium-pendulum training trials. C) Estimated peak cart speed averaged across the final four trials of the short-, medium-, and long-pendulum test conditions. D–F) Per-trial estimated peak cart speed during the 20 test trials with the short (D), medium (E), and long (F) pendulums. G–I) Corresponding per-trial estimated peak pendulum angular speeds for the short (G), medium (H), and long (I) pendulums. Lines show group means, shaded bands denote ±1 SE across participants, and error bars in C denote ±1 SE across participants. Peak speeds were defined as the maximum absolute estimated linear or angular velocity within each trial.

Estimated peak pendulum angular speed over the final four trials differed across pendulum conditions (Fig. 6G–I). Mean estimated peak angular speed was 231.817 ± 40.516 °s^−1^ for the short assembly, 164.558 ± 37.198 °s^−1^ for the medium assembly, and 100.314 ± 34.313 °s^−1^ for the long assembly. Mauchly’s test did not indicate a violation of sphericity, *W* = 0.585, *x*^2^(2) = 4.821, *p* = 0.090; therefore, uncorrected degrees of freedom were reported. The mixed repeated-measures ANOVA showed a significant main effect of pendulum condition on estimated peak angular speed, *F*(2, 20) = 47.549, *p* < 0.001, *ω*^2^ = 0.687. There was no significant main effect of presentation order, *F*(1, 10) = 0.564, *p* = 0.470, *ω*^2^ < 0.001, and no significant Condition × Order interaction, *F*(2, 20) = 1.336, *p* = 0.285, *ω*^2^ = 0.016. Holm-adjusted pairwise comparisons, averaged across presentation orders, showed that estimated peak angular speed was significantly greater for the short than for the medium assembly, for the short than for the long assembly, and for the medium than for the long assembly (all *p* < 0.001).

## Discussion

We examined human balancing of three physical linear-rail cart–pendulum assemblies whose passive pendulum dynamics were characterized empirically. Balance duration was greater in the final four than in the first four training trials with the medium assembly, whereas no detectable start-to-end change occurred within any subsequent test block. In late-block testing, the short assembly, which exhibited the fastest passive dynamics, was balanced for less time than the medium and long assemblies, which did not differ detectably from one another. This pattern was consistent with the prediction that the faster short assembly would be more difficult to balance. Together, the passive and human measurements provide a quantitative reference for future comparisons of human and autonomous control under matched mechanical and task constraints.

### Passive pendulum characterization

The free-decay model was used to quantify the passive rotational dynamics of the three pendulum assemblies. The fitted inertia-normalized gravitational coefficients were within 3.4–5.3% of the corresponding nominal values for uniform bare rods, supporting the physical plausibility of the fitted gravitational term. The damping and Coulomb-like friction coefficients should be interpreted as effective descriptive properties of the complete rod-and-mount assemblies. For the long assembly, the fitted Coulomb-like coefficient was substantially smaller than those of the other two assemblies and was therefore treated as effectively negligible or weakly resolved. Because only one free-decay recording was fitted for each assembly, these estimates characterize the tested apparatus rather than between-recording variability or parameter repeatability.

### Comparative results: passive mechanics and human control

The passive measurements revealed clear differences among the three assemblies. The short assembly exhibited the highest oscillation frequency and shortest passive-fall duration, whereas the medium and long assemblies had lower oscillation frequencies and longer passive-fall durations. These differences are consistent with different temporal margins being available for corrective action. Free-decay behavior reflected the combined effects of the fitted gravitational, viscous, and Coulomb-like terms and should therefore not be attributed to any single mechanical property, such as rod length or inertia.

In the active-balancing experiment, balance duration increased from the first four to the final four training trials with the medium assembly. During the subsequent test blocks, no statistically detectable start-to-end change occurred for any assembly. Late-block balance duration was shortest for the short assembly and was longer for the medium and long assemblies, with no statistically detectable difference between the latter two. This pattern is consistent with the familiar mechanical result that shorter inverted pendulums have faster passive dynamics and provide less time for corrective action than longer pendulums.

Because the medium condition served both as the training condition and as the first test condition for every participant, comparisons involving the medium assembly should be interpreted relative to a trained, fixed-order reference rather than as fully counterbalanced condition effects. The short–long comparison was less directly affected by this design feature because the order of those two conditions was counterbalanced. Nevertheless, medium-condition performance may partly reflect prior task-specific practice and its fixed position as the first test block.

Estimated peak cart speed and peak pendulum angular speed showed different condition-dependent patterns. Mean estimated peak cart speed was numerically highest in the medium condition and did not vary monotonically across assemblies, whereas estimated peak pendulum angular speed was highest for the short assembly and lowest for the long assembly. The short and medium conditions did not differ detectably in estimated peak cart speed. Thus, estimated peak cart speed did not mirror the pattern of balance duration across assemblies. Because these speed estimates were derived from approximately 6-Hz position records, they were treated as approximate descriptive measures, and no stronger conclusion about participants’ control strategies was drawn.

### Relation to previous work

Previous studies of human balancing have emphasized the roles of delayed feedback, passive mechanics, and intermittent corrective action [12–14,16,31]. The present findings complement this work by showing that the physical assembly with the fastest measured passive dynamics was balanced for less time than the two slower assemblies.

Time-delayed feedback is particularly consequential in an unstable system because corrective action is based on state information that is already out of date. The poorer performance observed with the faster short assembly is therefore consistent with a smaller effective margin for correction before threshold crossing. Intermittent or burst-like corrections have also been reported in stick-balancing and cart–pendulum tasks [12,16,31], although the present analysis did not test whether participants used such a strategy.

Sensorimotor delay, sensory and motor noise, and constraints on human corrective action were not measured directly and should therefore be regarded as candidate mechanisms rather than demonstrated explanations for the observed differences among assemblies. Because linear cart–pendulum systems are widely used as benchmark plants in control engineering, the paired passive and human measurements provide a quantitative reference for evaluating autonomous controllers under matched physical and task conditions. They may also support comparisons across controller architectures, sensory constraints, and participant populations.

### Limitations

Several limitations should be considered. The human sample was modest (n = 12), and the medium assembly served both as the training condition and as the first test condition. Only one free-decay trajectory was fitted for each assembly, passive falls were released manually by a single operator, and the assemblies differed in rod material, mass, and attachment mass as well as length. Balance duration was capped at 10 s, and the approximate speed estimates were derived from position records sampled at approximately 6 Hz. Although this sampling rate limits the precision of the derived kinematic measures, it is less consequential for the primary balance-duration outcome, which was determined from the timestamped angular trajectory as the interval from trial onset to the angular-threshold crossing or the 10-s trial limit. Finally, the free-decay model characterized fixed-pivot rotational dynamics rather than the complete coupled cart– pendulum dynamics present during human control. These factors limit the generalizability of the human findings, the assessment of mechanical parameter repeatability, and the precision of the kinematic interpretation.

### Future work

An important next step will be to compare human performance with that of a model-based controller using the same pendulum assemblies on a motorized version of the apparatus. Such a comparison could provide an engineered reference, helping to distinguish features of performance that arise from the task dynamics from those associated with biological sensorimotor constraints. A controller could be developed by combining the fitted rotational coefficients with independently characterized cart dynamics, pendulum–cart coupling, actuation properties, and rail constraints. Computational models based on our previous dual linear-quadratic regulatory framework [28,29] could then be extended to include sensory delay, sensory and motor noise, friction, and actuation limits.

Candidate controllers could first be evaluated in simulation and subsequently implemented on the same or a closely matched physical apparatus. Maintaining common pendulum dynamics, rail limits, initial conditions, and failure criteria would permit informative comparisons between human and autonomous balancing. This approach could test which aspects of the observed differences in balance duration and approximate movement kinematics follow from the mechanical and task constraints alone, and which are reproduced only when biologically relevant sensorimotor constraints are included.

## Author Contributions

LAH and ISH conceived and designed the study. LAH implemented the experimental protocol, collected the data, and performed the primary data processing and human-participant analyses. ISH reviewed and verified the human-participant analyses and developed and conducted the passive free-decay modeling. LAH wrote the initial draft of the manuscript. Both authors interpreted the results, reviewed and edited the manuscript, and approved the final version.

## Data Availability

The anonymised data supporting the findings of this study are publicly available in Zenodo at https://doi.org/10.5281/zenodo.21979469.

## Supplementary Information

**Table S1.** Parameter bounds and condition-specific initial values used for free-decay model fitting.

| Parameter | Units | Lower bound | Upper bound | Short initial | Medium initial | Long initial |
| --- | --- | --- | --- | --- | --- | --- |
| $k$ | $s^{-2}$ | 1.0 | 100.0 | 47.0 | 23.0 | 15.0 |
| $d$ | $s^{-1}$ | $1.0 \times 10^{-4}$ | 0.5 | 0.050 | 0.020 | 0.028 |
| $q$ | $s^{-2}$ | $1.0 \times 10^{-5}$ | 0.5 | 0.050 | 0.020 | 0.016 |

The smoothing velocity, *ω*_3_, was fixed at 0.0087 rad *s*^−1^(0.5° *s*^−1^) for all three assemblies and was not estimated during optimization. Parameter bounds and initial values are shown in physical units; *k, d*, and *q* were transformed into logarithmic coordinates for optimization.

**Table S2.** Optimization settings used for free-decay model fitting.

| Setting | Value |
| --- | --- |
| Optimizer | MATLAB lsqnonlin |
| Optimization algorithm | Trust-region-reflective |
| Model integration | MATLAB ode45 |
| Model evaluation times | Original measurement timestamps |
| Finite-difference type | Central |
| Finite-difference step size | $1.0 \times 10^{-3}$ |
| Maximum optimization starts | 5 |
| First starting point | Condition-specific initial values reported in Table S1 |
| Additional starting points | Random local perturbations around the condition-specific initial values in logarithmic parameter space |
| Standard deviation of logarithmic perturbations | 0.7 |
| Starting-point constraints | Perturbed starting points restricted to the specified parameter bounds |
| Selection criterion | Smallest residual sum of squares among successfully completed attempts, irrespective of optimization exit flag |
| Early-stopping criterion | Angular RMSE $\leq 0.5^\circ$ |
| Maximum iterations | 1000 |
| Maximum function evaluations | 10,000 |
| Function tolerance | $1.0 \times 10^{-10}$ |
| Step tolerance | $1.0 \times 10^{-10}$ |
| Optimality tolerance | $1.0 \times 10^{-8}$ |

